# Lysinoalanine crosslinking in the extracellular flagellar hook of Synergistota

**DOI:** 10.64898/2026.05.22.727228

**Authors:** Michael J. Lynch, Nyles W. Charon, Brian R. Crane

## Abstract

Bacterial motility is driven by the flagellar motor, a conserved macromolecular complex that enables cells to navigate chemical gradients. In spirochetes, motility depends on periplasmic flagella whose function is supported by a unique post-translational modification (PTM) in the hook protein FlgE: a covalent lysinoalanine (Lal) crosslink between adjacent subunits. This modification has been shown to be conserved across the phylum and is required for motility. In spirochetes, Lal is thought to provide additional hook stability needed to meet the demands of flagellar rotation in the confined space of the periplasm and associated load of the wrapped cell body. Guided by a bioinformatic screen based on conserved catalytic residues required for Lal formation, we investigated the possibility that bacteria with extracellular flagella also produce Lal. Using a combination of biochemical crosslinking assays and high-resolution mass spectrometry, we demonstrate that FlgE from *Thermanaerovibrio acidaminovorans,* a motile member of the Synergistota phylum, forms Lal crosslinks analogous to those of spirochetes *in vitro* and in wild type cell culture. These results establish the first example of Lal crosslinking in extracellular flagellar hooks and the first instance outside of the spirochete phylum. Thus, Lal crosslinking is more broadly conserved than previously appreciated and may represent a general mechanism for enhancing hook stability and function for extracellular as well as periplasmic flagella.

**Importance:** Many bacteria employ complex rotary engines known as flagella to propel them toward nutrients or away from harmful environments. In spirochetes, motility depends on lysinoalanine (Lal) crosslinks within the flagellar hook protein FlgE to strengthen the universal joint of the rotating filament, which resides in the periplasm. This study shows that the same type of modification also exists in a bacterium distantly related to spirochetes, *Thermanaerovibrio acidaminovorans*, that contains more typical extracellular flagella. Using biochemical experiments and mass spectrometry, we confirm that its flagellar hook protein self-catalyzes Lal crosslinks, marking the first time this modification has been observed outside spirochetes and in extracellular flagella. These findings suggest that Lal crosslinking is more widely conserved than previously thought and may represent a general strategy bacteria use to reinforce their flagella. Because motility is closely linked to survival and the ability of bacteria to cause disease, this discovery improves our understanding of bacterial motility mechanisms and how it is maintained across diverse species.

## Introduction

Flagellated bacteria move within their environment, propelled by the rotary action of membrane-embedded motors.^1,2^ Powered by the ion gradient spanning the inner membrane, counterclockwise and clockwise rotation of long filaments push the cell forward and allow the cell to change direction^3^. Rotational control over the bacterial flagellar motor (BFM) is achieved via interactions between the cytosolic portion of the BFM, known as the C-ring, and the phosphorylated form of the chemotaxis response regulator CheY.^3–6^ CheY is phosphorylated by the histidine kinase CheA, whose activity is modulated by chemoreceptor arrays in the inner membrane.^3,7,8^ Overall, this system allows bacteria to sense extracellular chemical gradients over several orders of magnitude and navigate towards nutrients or away from toxins.^3^

The BFM is a complex macromolecular assembly comprised of over 30 different proteins present in stoichiometries ranging from one to several thousand.^1,9,10^ Despite being a highly complex and dynamic nanomachine, the overall architecture of the BFM is well conserved (Figure 1A).^11^ In most species, a 5–20-micron helical filament extends into the extracellular space, attaching to the cell body via the hook. The hook is a ∼55 nm long tubular structure comprised of >120 subunits of a single protein, FlgE.^12–18^ Serving as a flexible adaptor, the hook converts the torque generated by the motor to the force used to propel the cell forward.^19^ The hook attaches to the rod, which serves as the drive shaft of the motor and is enclosed by several rings that span the outer membrane (L-ring), peptidoglycan layer (P-ring), and inner membrane (MS-ring).^14,20^ At the base of the motor is the C-ring, which, in addition to binding CheY-P and the membrane-embedded stator units, also interacts with the export apparatus machinery that controls protein secretion during motor assembly.^20–25^

**Figure 1:**
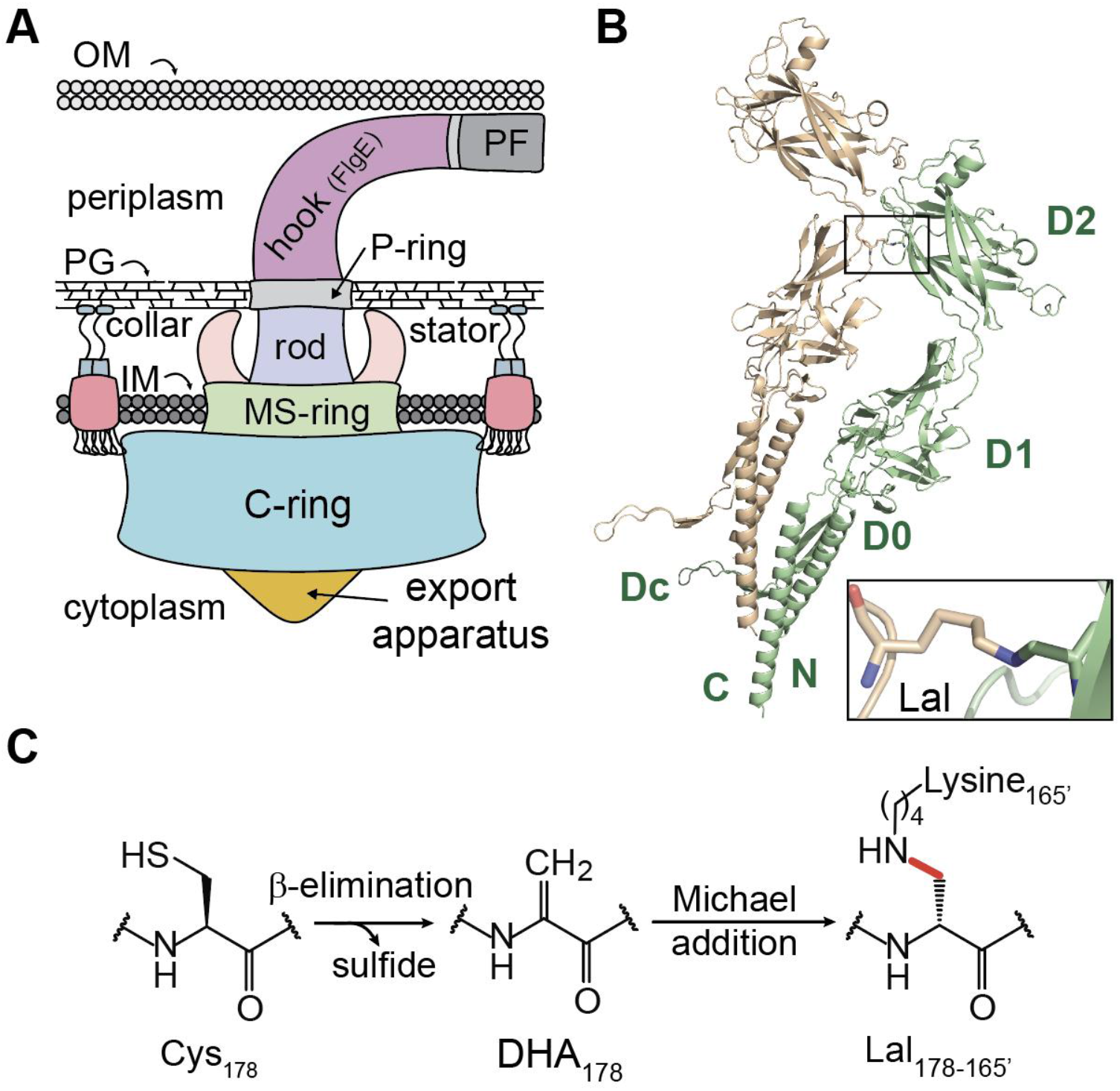
Overview of flagellar motor architecture and Lal crosslinking. A) General structure of the spirochete periplasmic flagellar motor. B) Structural model of a *T. denticola* FlgE dimer^31,84^ crosslinked via lysinoalanine (Lal, denoted with black box and shown in the lower right inset). C) Lal crosslink formation mechanism, wherein Cys178 undergoes a β-elimination to yield the dehydroalanine (DHA) intermediate and release sulfide. DHA then reacts with Lys165 from an adjacent FlgE monomer in an aza-Michael addition to yield the mature Lal crosslink.

Despite being well conserved, variations in flagellar motor architecture meet different environmental or ecological niches.^11,26^ One of the most striking differences is observed in members of the phylum Spirochaetota. Spirochetes are unique in that they enclose their flagellar filaments within the periplasm between the peptidoglycan layer and the outer membrane (Figure 1A).^27,28^ Attached to BFMs localized at the cell poles, filaments wrap around the cell and in some species form a continuous ribbon along the entire cell body.^28–30^ Filament rotation pushes and pulls on the cell cylinder, inducing undulations that propel the cell forward.^28,29^ We found that the flagellar hook protein FlgE catalyzes a unique post-translational modification (PTM) in the form of a lysinoalanine (Lal) crosslink between adjacent FlgE subunits in the hook (Figure 1B). ^31–33^ Lal formation in FlgE results in the formation of high-molecular weight complexes (HMWCs) in both anti-FlgE western blots of spirochete whole-cell lysate and from recombinant spirochete FlgE over-expressed and purified from *E. coli*.^31,32^ Importantly, these Lal crosslinked HMWCs are absent from non-spirochete species (ex. *Salmonella enterica*^34^), suggesting this PTM may be a specific adaptation to the phylum. The crosslink forms between a conserved cysteine and lysine residue in the FlgE D2 and D1 domains, respectively.^31^ Formation of Lal proceeds in two distinct biochemical steps. First, cysteine undergoes a β-elimination, releasing hydrogen sulfide and forming dehydroalanine (DHA).^31^ Next, the β-carbon of DHA reacts with the ε-nitrogen of lysine via an aza-Michael addition to form the mature Lal crosslink (Figure 1C).^31^ To date, we have confirmed Lal crosslinking in FlgE samples from six representative spirochete species from three out of the four orders across the phylum.^33^ These data suggest that Lal crosslinking is a conserved post-translational modification (PTM) across the phylum. Importantly, mutagenesis of catalytic or structural residues that abolish Lal formation *in vivo* resulted in normal hook and filament assembly, but rendered the cells non-motile in swimming assays.^32,33^ These findings are significant because motility is a key virulence determinant in many bacterial pathogens, including pathogenic spirochetes, which cause a range of human and animal diseases, such as Lyme disease, periodontal disease, syphilis, pinta, yaws, endemic syphilis, leptospirosis, swine dysentery, and bovine digital dermatitis.^35–44^ We have recently shown that Lal crosslinking can be targeted by small molecule inhibitors, which act on cells to curtail their motility and hence infectivity.^45,46^

We now extend these findings beyond bacteria with periplasmic flagella and demonstrate Lal crosslinking in the flagellar hook of the bacterium *Thermanaerovibrio acidaminovorans* (*T. acidaminovorans*, Ta). *T. acidaminovorans* is a Gram-negative, moderately thermophilic anaerobe originally isolated from a sugar refinery in the Netherlands in the 1990s.^47^ Interestingly, *T. acidaminovorans* cells are curved rods with multiple lateral flagella on the concave side of the cell body that extend outside of the cell body.^47,48^ *T. acidaminovorans* was identified by a bioinformatic screen, where we filtered all known annotated bacterial FlgE sequences for the catalytic residues required for Lal biogenesis. This search produced a list of potential Lal crosslinking candidates, with a majority of the hits being species from the Synergistota phylum, suggesting that Lal crosslinking may be a conserved feature amongst motile Synergistota species. The Synergistota phylum is presently divided into a single recognized class, Synergistia, and one order, Synergistales. These taxa encompass 22 known genera assigned to eight families, including Acetomicrobiaceae, Aminiphilaceae, Aminobacteriaceae, Dethiosulfovibrionaceae, Synergistaceae, Thermovirgaceae, as well as two unnamed families.^49–51^ Organisms within this phylum have been recovered from a broad range of anaerobic habitats, including wastewater treatment reactors, soils, animal and insect digestive systems and petroleum-associated environments^50,52^. Despite their environmental diversity, Synergistota spp., including *T. acidaminovorans*, share the trait of fermenting amino acids instead of carbohydrates.^50^ Herein, we employ a combination of biochemical crosslinking assays and high-resolution mass spectrometry to show that *T. acidaminovorans* FlgE forms Lal crosslinks *in vitro* and *in vivo* similar to those observed in spirochetes. This represents the first reported instance of Lal crosslinking in extracellular flagellar hooks and the first identified outside the spirochete phylum.

## Materials and Methods

### i. Identification of FlgE crosslinking candidates

To identify lysinoalanine (Lal) crosslinking candidates, all annotated FlgE genes in the bacterial kingdom, excluding spirochete species, were retrieved using Annotree.^53^ FlgE sequences were then filtered, excluding the sequences without a “CNL” or “SNL” (Cys/Ser, Asn, Leu) sequence motif. Sequences were aligned with a multiple sequence alignment was performed using clustal Omega.^54^ Sequences containing all three catalytic motifs previous identified to be required for Lal formation: (1) CNL/SNL (conserved Cys/Ser, Asn and Leu), (2) the catalytic lysine and (3) the catalytic threonine were selected for analysis.

### ii. T. acidaminovorans FlgE expression and purification

The full-length *T. acidaminovorans* (DSM 6589, ATCC 49978, Su883 - BacDive ID: 17780, Loc_01371, CP001818 1378228-1380138 (rev)) *flgE* gene was *E.coli* BL21(DE3) codon-optimized with an N-terminal His-Tag and ordered from Twist Biosciences in pet28a+ (Figure S1). Plasmids were transformed into BL21 (DE3) *E. coli* cells, grown in LB media containing 50 μg/ml kanamycin to an optical density (OD_600_) of ∼0.3–0.4 at 37 °C then lowered to 17 °C, grown to an OD_600_ of ∼0.6–0.8, induced with 0.5 mM IPTG and shaken for 12-18 hours. Cells were harvested via centrifugation at 4,000xg and stored at -20 °C.

To purify *T. acidaminovorans* FlgE, cells from 4 L of culture were thawed on wet ice for 30 minutes, resuspended in 60 mL of 20 mM Tris pH 7.5, 500 mM NaCl, 5 mM imidazole and lysed via sonication for 12 minutes at an intensity of 70% with 2 second on/off pulses. Cell lysate was clarified via centrifugation at 100,000xg for 45 minutes at 4°C and passed through a 5 mL Ni-NTA affinity column at a flow rate of 1-2 mL/min. Ni-NTA resin was washed with 5-10 CVs of 20 mM Tris pH 7.5, 500 mM NaCl, 25 mM imidazole and bound proteins eluted in 1.5 mL fractions with 20 mM Tris pH 7.5, 500 mM NaCl, 250 mM imidazole. Fractions were tested with Bradford and positive fractions were combined. To remove the N-terminal affinity and SUMO tags, FlgE samples were mixed with ∼1 mg of His_6_-MBP-uTEV3 (gifted from Alice Ting, addgene plasmid # 135464 ; http://n2t.net/addgene:135464 ; RRID:Addgene_135464)^55^ and dialyzed against 20 mM Tris pH 7.5, 150 mM NaCl, 0.5 mM EDTA, 1 mM DTT for 12-18 hours at 4°C. Cleaved FlgE was buffer exchanged into 20 mM Tris pH 7.5, 500 mM NaCl, 5 mM imidazole and then passed over 1-2 mL of Ni-NTA resin to remove His6-MBP-uTEV3 and SUMOylated FlgE. The flow-through was collected, concentrated to 5-10mL using a 10 kDa MWCO filter and purified via S200 (26/60) size-exclusion chromatography at a flow rate of 2.5 mL/min in 20 mM Tris pH 7.5, 150 mM NaCl. The monomer peak was collected, confirmed via SDS-PAGE, concentrated to ∼1 mL and flash frozen in 25 uL aliquots. Protein concentrations were measured via the BCA assay and reported as the average concentration ± the standard deviation of three technical replicates.

### iii. In vitro Lal crosslinking assays

Crosslinking assays were performed as described previously.^31^ Briefly, FlgE samples were mixed 1:1 with 2x crosslinking buffer containing 50 mM Tris pH 8.5, 300 mM NaCl, 2 M ammonium sulfate. For DTNB- and NEM-treated samples, a final concentration of 1 mM of each compound was included. Samples were incubated for 48 hours at 4°C, mixed with 4X Laemmli buffer and visualized by SDS-PAGE. For the *T. acidaminovorans in vitro* FlgE MS samples, the final protein concentration was 3.0 mg/mL. Silver stain samples were adjusted to a final concentration of 1 mg/mL and stained according to the manufacturer’s guidelines. Crosslinking band intensities were quantified using FIJI as described previously.^31,45,56^

### iv. SDS-PAGE and in-gel digestion of Lal crosslinked FlgE

Crosslinked recombinant FlgE samples were directly mixed 1:1 with 2× SDS-PAGE loading dye supplemented with 100 mM βME and heated at 90°C for 10 min. Denatured samples were then centrifuged briefly and electrophoresed on a 4–20% denaturing Tris-glycine SDS-PAGE gel. The gel was stained with Coomassie blue, destained, and washed extensively with 18.2 MΩ water. Multimer HMWC bands were excised and submitted for MS. Excised bands were sliced into ∼1-mm cubes and washed consecutively with 200 μL deionized water followed by 50 mM ammonium bicarbonate, 50% (v/v) acetonitrile, and finally 100% acetonitrile. The dehydrated gel pieces were reduced with 50 μL of 10 mM DTT in 100 mM ammonium bicarbonate for 1 h at 60°C, followed by alkylation with 50 μL of 55 mM iodoacetamide in 100 mM ammonium bicarbonate at room temperature in the dark for 45 min. Wash steps were repeated as described above. The gel was then dried and rehydrated with 100 μL trypsin at 10 ng/µL in 50 mM ammonium bicarbonate and incubated on ice for 30 min and then at 37°C for 18 h. Digestion was stopped by adding 30 µL 5% (v/v) formic acid. The digested peptides were extracted from the gel twice with 200 μL of 50% (v/v) acetonitrile containing 5% (v/v) formic acid and once with 200 μL of 75% (v/v) acetonitrile containing 5% (v/v) formic acid. Extractions from each sample were pooled together. The pooled sample was dried in SpeedVac SC110 (Thermo Savant, Milford, MA) to 200 μL to remove the acetonitrile (ACN) and then filtered with a 0.22-µm spin filter (Costar Spin-X from Corning) and dried to dryness in the speed vacuum. Each sample was reconstituted in 0.5% (v/v) formic acid prior to HPLC-MS/MS analysis.

### v. Identification of in vitro Lal crosslinks by nano-LC-MS/MS and data analysis

The tryptic digests were analyzed by nano-LC-MS/MS analysis at the Cornell Proteomics and Metabolomics Facility. The analysis was carried out using an Orbitrap Fusion Tribrid (Thermo Fisher Scientific, San Jose, CA) mass spectrometer equipped with a nanospray Flex Ion Source and coupled with a Dionex UltiMate 3000 RSLCnano system (Thermo, Sunnyvale, CA). Each sample was loaded onto a nano-Viper PepMap C18 trapping column (5 µm, 100 µm × 20 mm, 100 Å, Thermo Fisher Scientific) at 20 µL/min flow rate for rapid sample loading. After 3 min, the valve switched to allow peptides to be separated on an Acclaim PepMap C18 nanocolumn (2 µm, 75 µm × 25 cm, Thermo Fisher Scientific) at 35°C in either 60- or 90-min gradients of 5 to 35% buffer B (98% ACN with 0.1% formic acid) at 300 nL/min (buffer A: 98% water, 2% CAN with 0.1% formic acid). The Orbitrap Fusion was operated in positive ion mode with nanospray voltage set at 1.85 kV and source temperature at 275°C. External calibrations for Fourier transform, ion trap, and quadrupole mass analyzers were performed prior to the analysis. Samples were analyzed using the CID and ETD toggle workflow, in which MS scan range was set to 350–1600 m/z, and the resolution was set to 120,000. Precursor ions with charge states of 3–7 were selected for ETD MS/MS acquisitions in ion trap analyzer and an automatic gain control (AGC) target of 3 × 10 (4). The precursor isolation width was 3 m/z, and the maximum injection time was 118 ms. Precursor ions with charge states of 2–3 were selected for CID MS/MS, and normalized collision energy was set to 30%. All data were acquired under Xcalibur 4.4 operation software and Orbitrap Fusion Tune application v3.5 (Thermo Fisher Scientific). All MS and CID-ETD MS/MS raw spectra from each sample were inspected manually using Xcalibur Qual Browser. Confident identification of Lal crosslinking peptides was achieved by manual confirmation of MS precursor ions with high mass accuracy and their associated CID and/or ETD MS/MS fragments using a peptide mass tolerance was 10 ppm and MS/MS fragment mass tolerance of 0.6 Da.

### vi. Site-directed mutagenesis of T. acidaminovorans FlgE

Site-directed mutagenesis was carried out using the QuikChange method and confirmed by DNA sequencing. The plasmid that expresses FlgE recombinant protein was used as a template to replace Cys180, Thr507 and Lys167 with alanine using primers MJL483/484, 489/490 and 495/496, respectively (see Table S1).

### vii. αFlgE western blot of T. acidaminovorans cell lysate

*T. acidaminovorans* (DSM 6589) cells from a 10 mL stationary-phase anaerobic culture were purchased from DSMZ GmbH and thawed on ice for 10 minutes. As a positive control, infectious *B. burgdorferi* B31 A3-68 (WT) cells (generously provided by Dr. Chunhao Li, Virginia Commonwealth University) were cultured in 10 mL of BSK-H Complete Medium (B8291) under microaerophilic conditions at 34°C for 4 days to stationary phase (∼10⁸ cells/mL). *B. burgdorferi* (Bb) cultures were harvested by centrifugation at 1,000 x g and stored at −80°C until use. Both *T. acidaminovorans* (Ta) and Bb cell pellets were thawed on ice for 10 minutes, washed with 200 μL of 18 MΩ water, and resuspended in 200 μL of 8 M urea. Samples were incubated with gentle agitation for 30 minutes at room temperature, and total protein concentrations were determined using a Bradford assay. As a negative control, 50 ng of recombinant Bb FlgE, produced in E. coli as previously described,^33^ was included. All samples were mixed 1:1 with 3.6x Laemmli sample buffer supplemented with 10% β-mercaptoethanol (βME), heated at 60°C for 5 minutes, and 10 μL of each sample was loaded onto a 50:50 4%/8% SDS-PAGE gel. Proteins were separated by electrophoresis at 200 V for 45 minutes, transferred to a PVDF membrane at 25 V for 16 hours, and blocked with 5% (w/v) skim milk in TBST (20 mM Tris-HCl, pH 7.5, 150 mM NaCl, 0.1% Tween 20) for 1 hour at 25°C. Membranes were incubated for 1 hour at 25°C with 10 mL of rabbit anti-Bb FlgE monoclonal IgG^57^ diluted 1:1,000 in 5% milk/TBST, washed three times with 25 mL TBST for 10 minutes each, and then incubated for 1 hour at 25°C with 10 mL of HRP-conjugated anti-rabbit IgG secondary antibody (Cell Signaling Technology, #7074) diluted 1:1,000 in 5% milk/TBST. Membranes were washed three additional times with 25 mL TBST for 10 minutes each before chemiluminescent detection using 1 mL of SuperSignal West Pico PLUS substrate and imaging on a ChemiDoc system

### viii. Identification of in vivo Lal crosslinks by nano-LC-MS/MS and data analysis

*T. acidaminovorans* (DSM 6589) cells purchased from DSMZ GmbH were washed with 200 μL of 0.22 μm-filtered Optima water and briefly centrifuged at 13,000 rpm. The resulting cell pellet was resuspended in 200 μL of 8 M urea, gently vortexed, and incubated at room temperature for 3 minutes. Cell lysates were then mixed 1:1 with 10% SDS, sonicated for 15 minutes, and centrifuged at 13,000 rpm. Approximately 13 μg of total protein lysate was loaded onto a 4–20% TGX Tris-glycine SDS-PAGE gel, separated by electrophoresis at 200 V for 30 minutes, and stained with Coomassie Blue. The upper portion of the stacking gel was excised, and in-gel reduction, alkylation, and tryptic digestion were performed as previously described. Tryptic peptides were analyzed as described above using electron-transfer/higher-energy collision dissociation (EThcD). All MS and EThcD MS/MS raw spectra from each sample were manually inspected using Xcalibur Qual Browser.

## Results

### i. T. acidaminovorans FlgE is a Lal crosslinking candidate

Using *T. denticola* FlgE as a query sequence and the three sequence motifs: (1) CNL (Cys, Asn, Leu), (2) catalytic lysine, and (3) threonine, we were able to identify a total of 175 FlgE sequences predicted to form lysinoalanine (Figure 2A-B). These sequences belong to a diverse group of bacterial species, including sequences derived from metagenomic data, uncultured samples, and/or environmental samples. Species belonging to the phylum Synergistota comprised most hits in our search (96/175 or ∼55%, Figure 2B), suggesting that Lal crosslinking may be a conserved PTM within the phylum. To reduce the number of potential candidates to test, we only considered FlgE sequences belonging to verified type-strain species that have been cultured in a clinical or laboratory setting. Based on these criteria, *T. acidaminovorans* was identified and selected as representative species for direct testing *in vitro* (Figure 2B-C).

**Figure 2:**
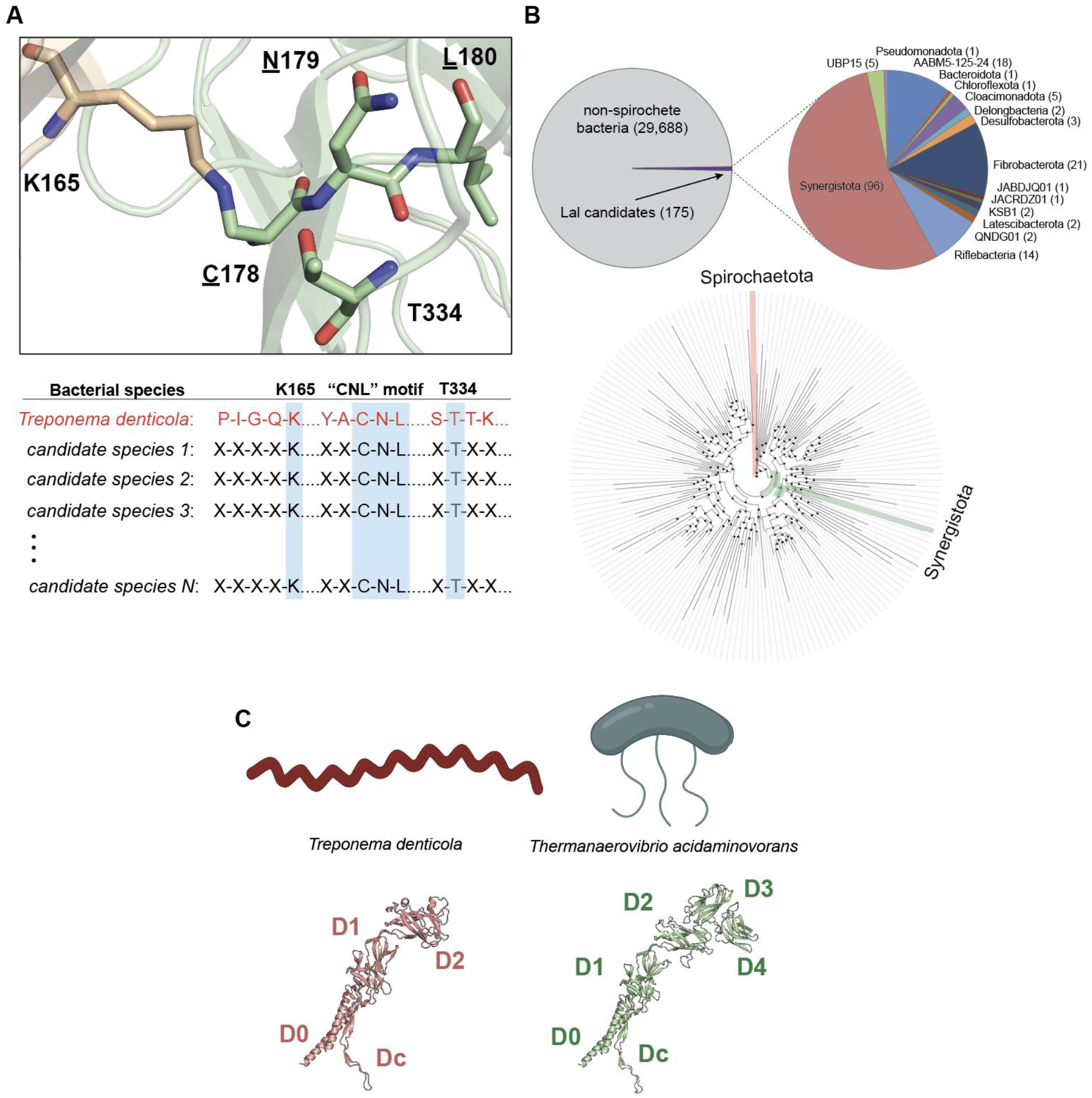
Identification of *T. acidaminovorans* as a FlgE Lal crosslinking candidate. A) (top) Location of the three sequence motifs in the FlgE Lal dimer structure used as search criteria in our bioinformatic screen. (bottom) FlgE sequence search strategy using a motif containing K165, CNL (<u>C</u>178, <u>N</u>179, <u>L</u>180) and T334 (Td numbering). B) Summary of Annotree hits following sequence motif filtering and filtering out spirochete FlgE sequences. (top) From the ∼30,000 sequences, 175 contained all three motifs, with 96 belonging to members of the Synergistota phylum. (B) Circular taxonomic tree at the phylum level with the Spirochaetota and Synergistota phylum highlighted. C) Comparison of cell shape (top) and FlgE structure (bottom) between *T. denticola* and *T. acidaminovorans*.

*T. acidaminovorans* is a Gram-negative, moderately thermophilic, strictly anaerobic bacterium whose cells are curved rods with multiple lateral flagella on the concave side of the cell body.^47^ Unlike spirochetes, *T. acidaminovorans* have external flagellar filaments.^47^ AlphaFold3^58^ models of the flagellar hook protein FlgE from *T. denticola* and *T. acidaminovorans* suggest that both FlgE protomers adopt a similar tertiary structure (Figure 2C), however, Ta FlgE is larger with two additional domains (D3 and D4) compared to Td FlgE.

### ii. T. acidaminovorans FlgE forms HMWCs in SDS-PAGE crosslinking assays

To test for Lal formation, we expressed, purified and tested *T. acidaminovorans* FlgE using *in vitro* Lal crosslinking assay, where we visualize high-molecular weight complex (HMWC) formation via SDS-PAGE. As shown in Figure 3A, when Ta FlgE is incubated in crosslinking buffer for 48 hours, robust HMWC formation is observed, with protein bands at molecular weight increments consistent with Lal crosslink formation. Addition of the cysteine-reactive compounds N-ethylmaleimide (NEM) and 5,5’-dithiobis-(2-nitrobenzoic acid) (DTNB) accelerates HMWC formation by reacting with Cys180 to favor dehydroalanine (DHA) formation, consistent with Lal crosslink formation preceding via cysteine conversion to DHA.^31^ Mutagenesis of the three catalytic residues Lys167, Cys180, and Thr507, previously identified in *T. denticola* FlgE,^31^ arrested HMWC formation at varying extends. Complete inhibition was observed for the C180A mutant, while the K167A mutant had small HMWC formation (Figure S2). Interestingly, the T507A mutant had reduced HMWC formation by ∼40% but was not completely crosslinking deficient (Figure S2). This result suggests that, unlike *T. denticola* FlgE, *T. acidaminovorans* FlgE Lal formation does not absolutely require the hydrogen bond supplied by the β-hydroxyl group of T507 for catalysis. To better visualize minor HMWC formation in the *T. acidaminovorans* FlgE mutants, *in vitro* SDS-PAGE assays were also performed in the presence of NEM and DTNB and silver-stained (Figure S2C). Silver staining revealed the presence of faint, non-specific protein bands that increased over time for the C180A mutant but were not dependent on the presence of NEM or DTNB. These bands were also present for the K167A mutant, but at higher levels, and increased upon the addition of NEM and DTNB. These bands are likely a result of non-specific FlgE-FlgE interactions that trigger Lal crosslink formation between DHA and other lysine residues in Ta FlgE.

**Figure 3:**
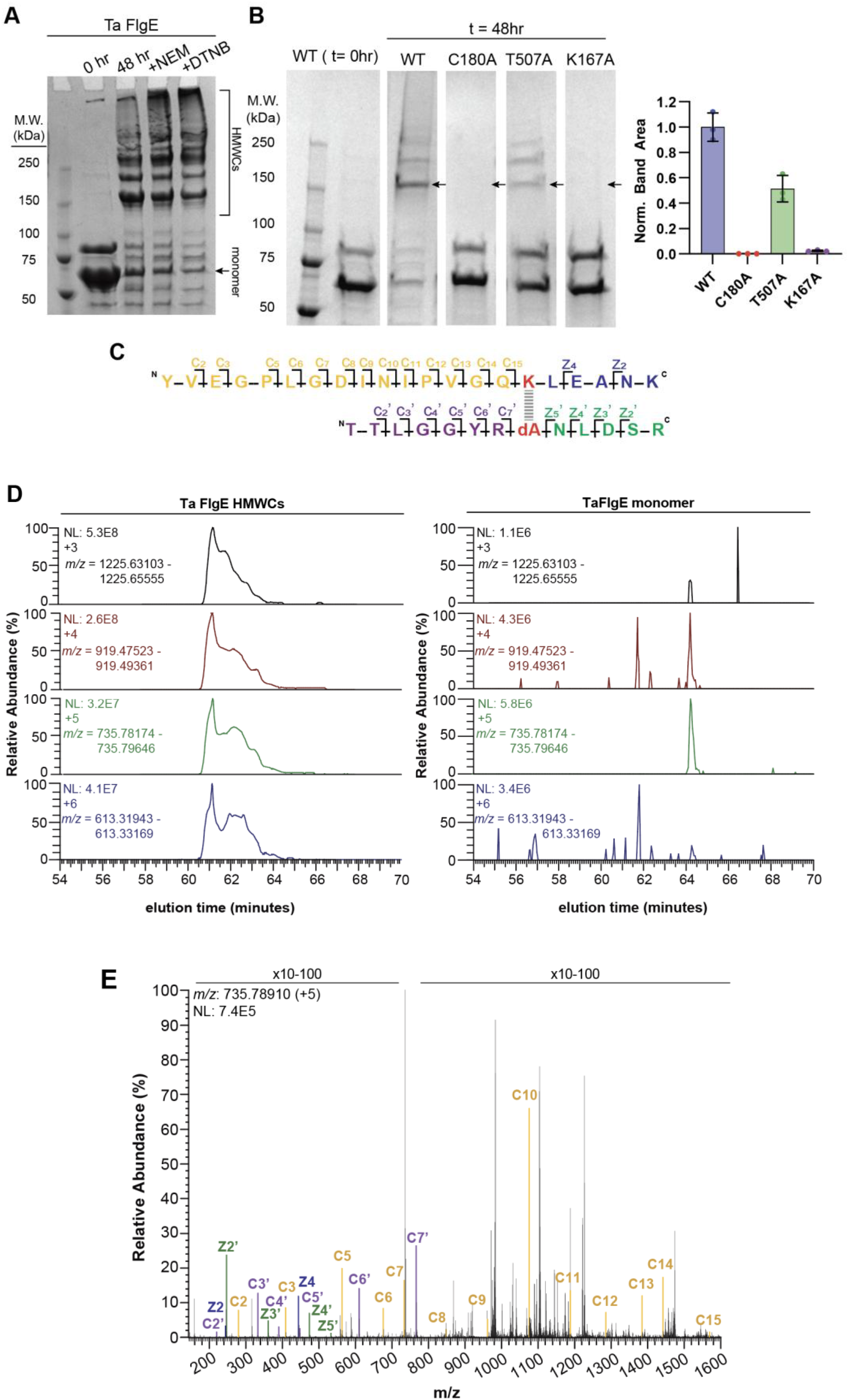
Detection of *T. acidaminovorans* recombinant FlgE Lal crosslinking peptides by LC-MS/MS. A) Coomassie-stained SDS-PAGE gel of *in vitro* Ta FlgE crosslinking assay. HMWC formation was assed at the 0 h and 48 h timepoints and with the addition of Td FlgE Lal crosslinking activators NEM and DTNB. Ta FlgE monomer and HMWC samples were submitted for Lal peptide detection via LC-MS/MS. B) (left) SDS-PAGE crosslinking assay of WT FlgE compared to catalytic alanine mutants C180A, T507A and K167A. For this assay, samples were run in triplicate (gel shown in Figure S2A) and the dimer band (denoted with black arrow) quantified (right). C) Primary structure of Ta FlgE Lal crosslinked peptide with c- and z-ions labeled. D) XICs of Lal crosslinked peptide m/z ranges in the [M+3H]^3+^ to [M+6H]^6+^ states from the HMWC (left) and monomer (right) datasets. E) ETD MS/MS fragmentation spectrum of [M+5H]^5+^ parent Lal peptide with c- and z-ions labeled as shown in (B).

### iii. Confirmation of Lal crosslinked peptides in T. acidaminovorans FlgE

To confirm that the HMWCs in Figure 2A are formed via Lal, we used LC-MS/MS to detect the peptides directly. Following separation by RP-HPLC, we detected the Lal-crosslinked peptide in two forms (Figure 3B-D and Figure S3). The two peptides are similar except for a missed cleavage at K176 position (denoted with *) in the peptide ^N^YVEGPLGDINIPVGQK*LEANK^C^ following trypsin digestion (Figure 3B). Missed cleavage at this position is not unexpected because Lal crosslinking likely sterically blocks cleavage by trypsin. As a result, the larger Lal crosslinked peptide is the dominant form detected in our data and is present in the [M+2H]^2+^ to [M+6H]^6+^ charge states (Figure 3C, left). The smaller Lal crosslinked peptide was also detected in the [M+3H]^3+^ to [M+5H]^5+^ charge states but was less abundant and produced fewer c- and z-ions compared to the higher molecular weight Lal peptide (Figure S3). Importantly, Lal peptide peaks are absent from the extracted ion chromatograms (XICs) from the Ta FlgE monomer sample (Figure 3C, right). Electron-transfer dissociation (ETD) MS/MS fragmentation of the [M+5H]^5+^ peptide produced c- and z-ions (sequence-specific peptide fragment-ion types produced by ETD) that allowed for the unambiguous assignment of the crosslinked peptide (Figure 3D and Figure S3). The parent ^N^C(CAM)NLDSR^C^ and ^N^YVEGPLGDINIVGQK^C^ peptides were also detected in multiple [M+XH]^x+^ charge states, as well as two forms of the DHA-containing peptide ^N^(DHA)^N^LDSRC (unmodified and βME-adduct; Figure S4). Collision-induced dissociation (CID) fragmentation of these peptides produced y- and b-ions consistent with each peptide, thereby confirming that Lal crosslink formation occurs via the DHA intermediate similar to Td FlgE Lal formation.^31^ Overall, these data confirm Lal crosslinking in *T. acidaminovorans* FlgE and represent the first reported case of Lal crosslinking outside of the spirochete phylum.

### iv. Detection of Lal crosslinked FlgE in T. acidaminovorans

To confirm that lysinoalanine (Lal) crosslinking occurs in *T. acidaminovorans* cells, we sought to detect Lal-crosslinked FlgE high-molecular-weight complexes (HMWCs) in *T. acidaminovorans* cell lysates by western blotting and mass spectrometry. *T. acidaminovorans* (DSM 6589) cells from a 10 mL stationary-phase anaerobic culture were obtained from DSMZ GmbH, and whole-cell lysates were separated by SDS-PAGE and analyzed by western blot using an antibody raised against *B. burgdorferi* FlgE. Conveniently, our *B. burgdorferi* FlgE antibody cross-reacted with *T. acidaminovorans* FlgE, albeit with substantially lower affinity, yielding a limit of detection of approximately 3 ng for Ta FlgE compared with <0.8 ng for Bb FlgE (Figure S5). Nevertheless, prominent FlgE-containing HMWCs were detected in *T. acidaminovorans* lysates (Figure 4A); 50 ng of monomeric recombinant Bb FlgE and Bb cell lysate as negative and positive controls for crosslinking, respectively. Compared with Bb cell lysate, Ta Lal-crosslinked hook complexes also migrated in the stacking portion of the SDS-PAGE gel, suggesting that *T. acidaminovorans* indeed catalyze the formation of Lal crosslinks within its flagellar hooks. We then excised the gel bands and subjected the HMWCs to HPLC-MS analysis, as described for the *in vitro* samples. Extracted ion chromatograms (XICs) revealed the presence of Lal-crosslinked FlgE peptides in both the [M+4H]⁴⁺ and [M+6H]⁶⁺ charge states, with retention times comparable to those observed for the *in vitro* samples (Figure 4B).

**Figure 4:**
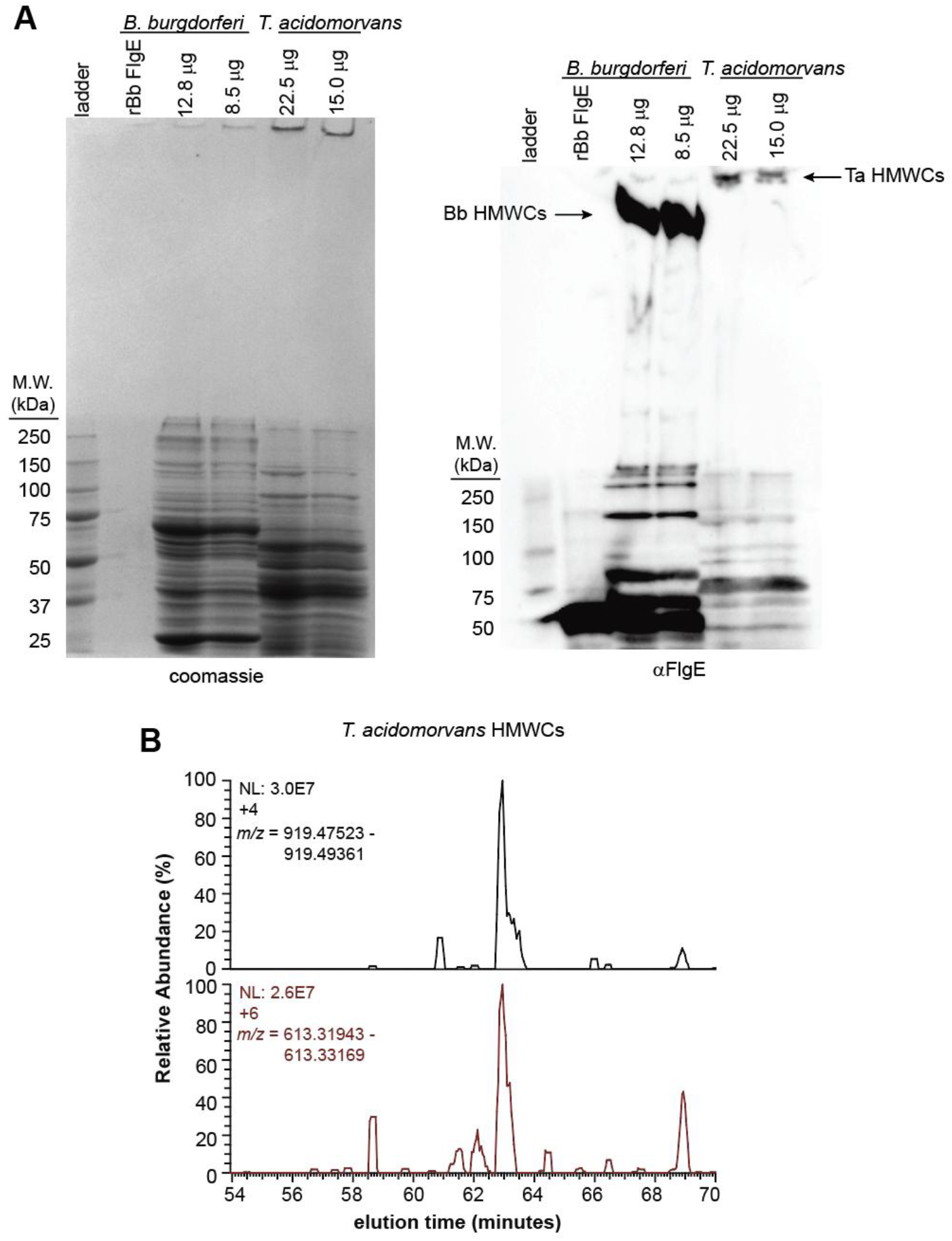
Detection from *T. acidaminovorans* cells of FlgE Lal crosslinked peptides by western blot and LC-MS/MS. A) Coomassie (left) and anti-FlgE western blot (right) of *T. acidaminovorans* whole cell lysate. Recombinant Bb FlgE and Bb whole cell lysate are included as a non-crosslinking and crosslinking control. B) XICs of Lal crosslinked peptide m/z ranges in the [M+4H]^4+^ and [M+6H]^6+^ states from the HMWCs present at the top of the coomassie and blot shown in (A). See Fig. 2 for comparison masses of *in vitro* crosslinked peptides.

### v. SNL-containing sequences expand the list of Lal candidates

In the order Leptospirales, bacterial species lack the catalytic cysteine in the CNL sequence motif and instead encode for a serine. Despite this difference, Lal crosslinking can still be detected in *Leptospira interrogans* FlgE *in vivo* from purified hooks or *in vitro* from recombinantly produced and crosslinked FlgE.^33^ Moreover, the cysteine to serine mutant in *T. denticola* FlgE can also readily undergo Lal crosslinking *in vitro* in SDS-PAGE assays and confirmed via HPLC-MSMS.^31^ Taken together, these data suggest that candidates containing a SNL motif should also be considered as Lal candidates if they also possess the two other search criteria. To provide a sense of the scope of such occurrences, we searched the collection of FlgE sequences according to the procedure previously described, yielding a total of 282 sequences that contain (1) a SNL motif, (2) catalytic lysine in the anticipated D1D2 linker region of FlgE and (3) a catalytic threonine (Figure S7). The species containing such FlgE proteins belonged to a diverse array of phyla, with Psuedomonadota and Desulfobacterota being the most represented members in the dataset.

## Discussion

Herein, we show that the flagellar hook protein from the Synergistota species *T. acidaminovorans* self-catalyzes lysinoalanine (Lal) crosslinks. Through a combination of biochemical, site-directed mutagenesis, and mass spectrometry experiments, we show that Lal crosslinking occurs through a mechanism similar to that of spirochete FlgE, including the pathogenic species *Treponema denticola*, *Borrelia burgdorferi*, *Treponema pallidum*, and *Leptospira interrogans*. In all cases, Lal crosslinking proceeds via the conversion of cysteine to DHA, followed by an aza-Michael addition involving the active-site lysine (Figure 1C).

Interestingly, one notable difference between Lal crosslinking mechanisms in spirochete and *T. acidaminovorans* FlgE is the dispensability of the hydrogen-bonding interaction between T334/507 (Td/Ta numbering) in Ta FlgE compared to Td (Figure S2). In Td FlgE, T334 was shown to be essential for Lal formation, with the T334A substitution completely abolishing Lal formation. Mass spectrometry and crystallographic data revealed that T334 is required for Lal formation because of its role in activating DHA for aza-Michael addition.^31^ Specifically, the T334 side-chain hydroxyl group forms an important hydrogen bond with the backbone amide nitrogen of N179, increasing the electrophilicity of the Cβ carbon of DHA and priming it for reaction with the ε-nitrogen of K165. The Ta FlgE T507A mutation results in an approximately 40% reduction in HMWC formation compared to WT FlgE (Figure S2), suggesting that it plays a similar role but is not strictly necessary for DHA activation and Lal formation.

One question that arises is exactly why *T. acidaminovorans* would require Lal crosslinking within its flagellar hook. Currently, Lal crosslinking in spirochete FlgE is believed to have evolved because of the higher hook strength required due to the unique motility of spirochetes. However, unlike spirochetes, *T. acidaminovorans* has external flagella, thus it remains unclear why *T. acidaminovorans* would catalyze Lal crosslinks between its FlgE subunits. One hypothesis is that it may be due to the unique shape of the *T. acidaminovorans* cells and the location of the flagella. *T. acidaminovorans* cells are shaped like curved rods with the lateral flagella localized to the concave side of cell body.^47,48^ This combination of cell shape and flagellar arrangement is rare and appears to be unique to motile members of the Synergistota. Curved morphologies are found in bacteria such as *Vibrio cholerae, Caulobacter crescentus*, and *Helicobacter pylori* and play an important role in the motility, and in some cases, pathogenicity of these species.^59–63^ For example, the helical morphology of *Helicobacter pylori* is thought to facilitate motility through the viscous gastric mucus by enabling a corkscrew-like mode of movement,^60,61^ while the curvature of *C. crescentus* has been shown to provide an evolutionary advantage by enhancing surface colonization in flowing environments.^63^ However, these organisms typically possess single or multiple polar flagella rather than laterally localized systems. Furthermore, lateral flagellar are observed in species such as *Vibrio parahaemolyticus, Vibrio alginolyticus and Shewanella putrefacient*, but often function as secondary flagellar systems, working in conjunction with polar flagella.^64–66^ In these species, the lateral flagellar system is thought to provide superior performance for swarming or for swimming under viscous conditions.^64–66^ Therefore, although the motility characteristics of *T. acidaminovorans* have not been quantified, it is possible that localization of their flagella on the face opposite a large convex surface demands high flagellar torque, and therefore stronger hooks, to translate the cell.

In support of this hypothesis, *T. acidaminovorans* FlgE contains two additional structural domains, D3 and D4, relative to *T. denticola* FlgE (Figure 2C). The presence of extra FlgE domains in *Campylobacter jejuni* and several related bacterial species enhance hook stability and robustness by increasing protein–protein interactions within the hook structure.^18^ Consistent with this role, Cj FlgE mutants lacking the D3 and D4 domains exhibited markedly reduced motility compared with WT cells, and light microscopy revealed that approximately 90% of cells possessed broken flagellar filaments.^18^ Since *T. acidaminovorans* FlgE also contains D3 and D4 domains, this suggests these bacteria may also require hooks with increased strength and stability.

*T. acidaminovorans* was selected for *in vitro* and *in vivo* testing as the representative species from the top phylum identified in our bioinformatic screen (Figure 2A–B). To gain insight into how well Lal crosslinking is conserved across the phylum, we mapped the Synergistota species from our screen onto a taxonomic tree generated using near-full-length 16S rRNA gene sequences adapted from McSweeney *et. al.* (Figure S6).^50^ Motile and non-motile species are both present within the Synergistota phylum; however, all motile species are predicted to encode FlgE proteins capable of forming Lal crosslinks. Interestingly, within the motile Syngeristota species outlined in Figure S6, there is some variability in flagellar location. Whereas most motile species have lateral flagella (ex. *Aminobacterium mobile,*^67^ *Dethiosulfovibrio acidaminovorans,*^68^ *Dethiosulfovibrio peptidovorans*^69^), some species possess only polar flagella (*Thermovirga lienii*^70^ and *Acetomicrobium hydrogeniformans*^71^), both polar and lateral flagella (ex. *Acetomicrobium flavidum*^72^) or are peritrichously flagellated (ex. *Aminiphilus circumsciptus*^73^). Given the diversity of habitats occupied by Syngeristota species, along with their distinctive cell morphology and flagellar organization, Lal crosslinking is more likely an adaptation related to cell structure than to environmental conditions. Consistent with this, the available data suggest that, as in the spirochete phylum, Lal crosslinking represents a conserved post-translational modification throughout Synergistota. Importantly, this is the first reported instance of Lal crosslinking in extracellular flagellar hooks and the first identified outside of the spirochetes.

Lastly, it should be noted that in our initial bioinformatic screen, we only considered sequences containing a conserved cysteine, rather than a serine, as the catalytic residue. This was done to reduce the pool of potential candidates and increase the probability of detecting Lal crosslinking in diverse FlgE orthologs, since cysteine is more reactive than serine in spirochete FlgE.^31^ However, within the spirochete phylum, *Leptospira* spp. possess a unique Lal crosslinking mechanism compared to other, more distantly related spirochetes.^33^ Specifically, they contain a conserved serine as the catalytic residue, which presumably eliminates hydroxide or water to form DHA, and can potentially form different Lal crosslinks between DHA and multiple conserved lysine residues.^33^ To incorporate these differences into our screen, non-spirochete FlgE sequences were re-screened and retained if they contained a conserved “SNL” motif (Figure S7). Using an approach similar to that of our initial screen, we identified over 282 candidates, the majority of which belonged to the Pseudomonadota phylum (193 sequences, or 68%). Although additional *in vitro* and *in vivo* testing will be required to confirm Lal crosslinking in these species, notable candidates include the BSL2 organisms *Shewanella algae* and *Shewanella putrefaciens*; the fish pathogens *Shewanella oncorhynchi* and *Pseudomonas pisciculturae*; and the fungal or plant pathogens *Pseudomonas tolaasii*, *Pseudomonas marginalis*, *Pseudomonas viridiflava*, *Pseudomonas salomonii*, *Pseudomonas lactucae*, and *Pseudomonas cyclaminis*.^74–82^ Moreover, *Leptospira* spp. and the candidate SNL species are not the only examples of variations in crosslinking that have been reported. In some contexts, cysteine and histidine residues have also been observed to react with DHA to form different types of inter-protein crosslinks.^83^ We found no examples of His and Cys substituting for Lys in the most conserved position of the CNL containing FlgE proteins; however, the position of the nucleophilic residue relative to the CNL/SNL motif can be variable, as it is in a loop region, and thus it is possible that non-Lys residues, at non-standard positions could be involved in a DHA-mediated linkage. Overall, Lal crosslinking in the flagellar hook protein FlgE, initially thought to be specific to the spirochete phylum, may represent a more widespread post-translational modification than previously appreciated.

## Author Contributions

M.J.L., N.W.C. and B.R.C. conceived the study. M.J.L. prepared samples, ran experiments, analyzed data and prepared figures. M.J.L. and B.R.C. wrote and edited the manuscript. B.R.C. supervised the study.

## Acknowledgements

We thank the Proteomics and Metabolomics Facility of Cornell University for providing the mass spectrometry data and NIH SIG grant 1S10 OD017992-01 support for the Orbitrap Fusion mass spectrometer. We also would like to thank Elizabeth Anderson, Qin Fu, and Sheng Zhang for their help with MS sample preparation and assistance in running the samples.

## Data Availability

All mass spectrometry data have been deposited to the ProteomeXchange Consortium via the PRIDE partner repository with the identifier PXD084241. All other data are included in the article and/or supporting information.

## Supplemental Information

**Table S1:**
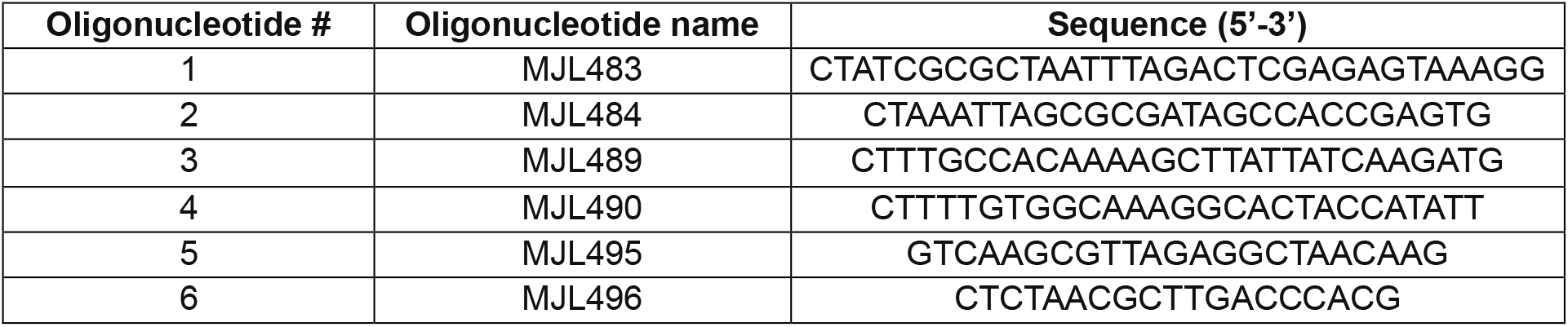
Primers used in this study.

**Figure S1:**
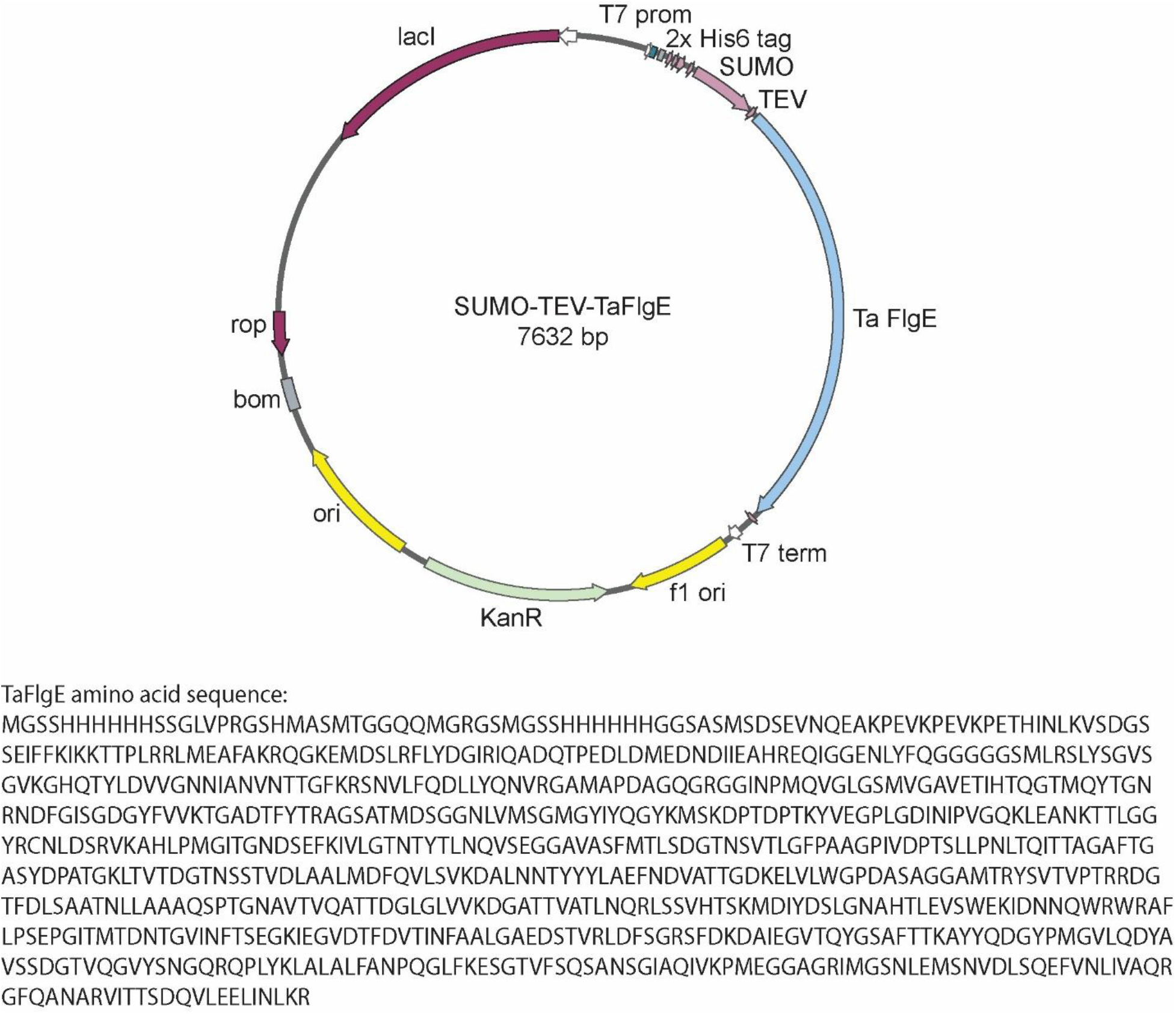
*T. acidaminovorans* FlgE overexpression vector map and amino acid sequence.

**Figure S2:**
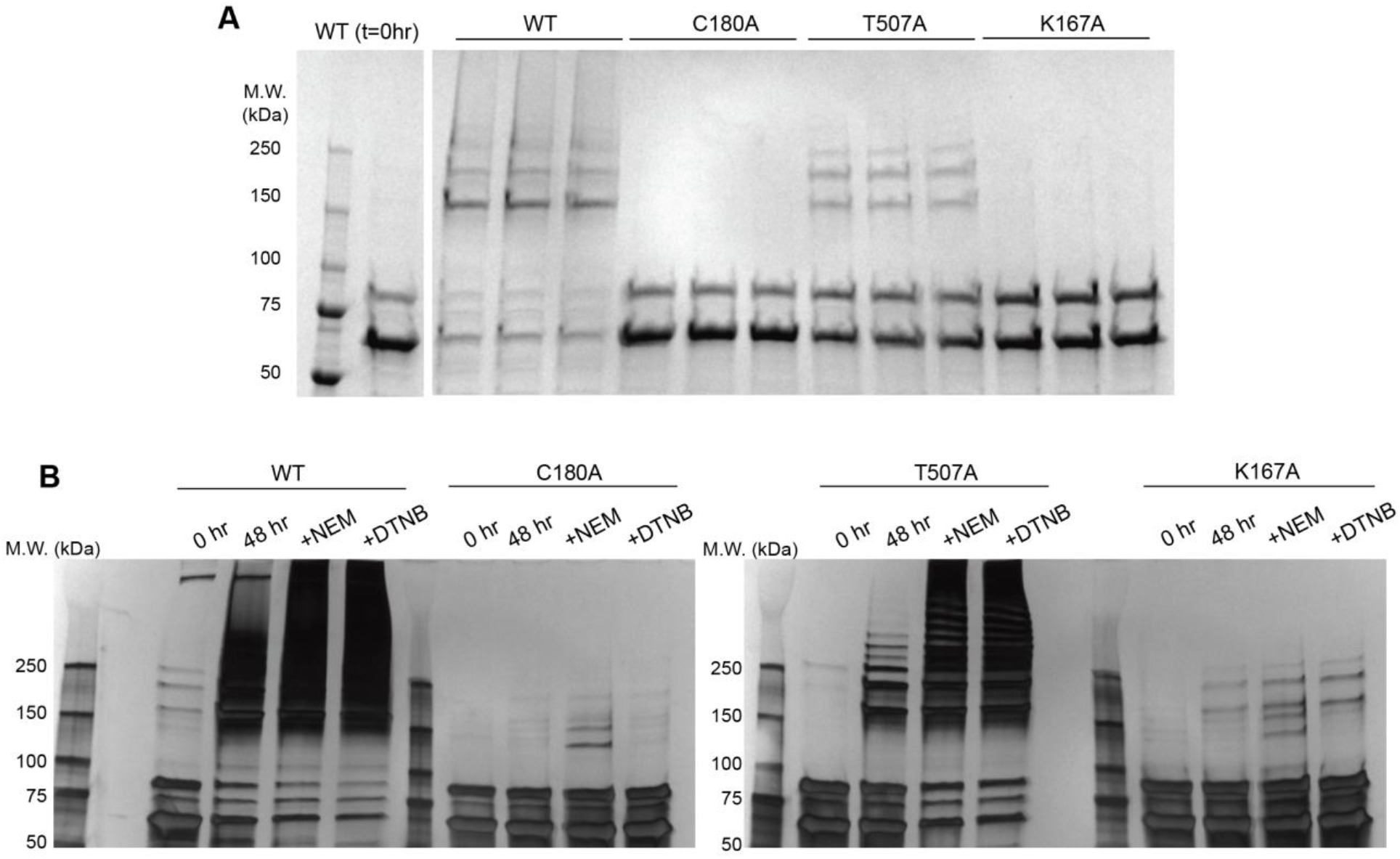
*In vitro* Lal crosslinking assays of *T. acidaminovorans* FlgE catalytic mutants. A) Coomassie-stained SDS-PAGE gels of WT, C180A, T507A and K167A mutant 48 h timepoint Lal crosslinking samples in triplicate. B) Samples down in (A) but stained with silver stain to visualize fainter HMWC bands. In addition to 48 h timepoint, 0 h and 48 h +NEM or +DTNB are included.

**Figure S3:**
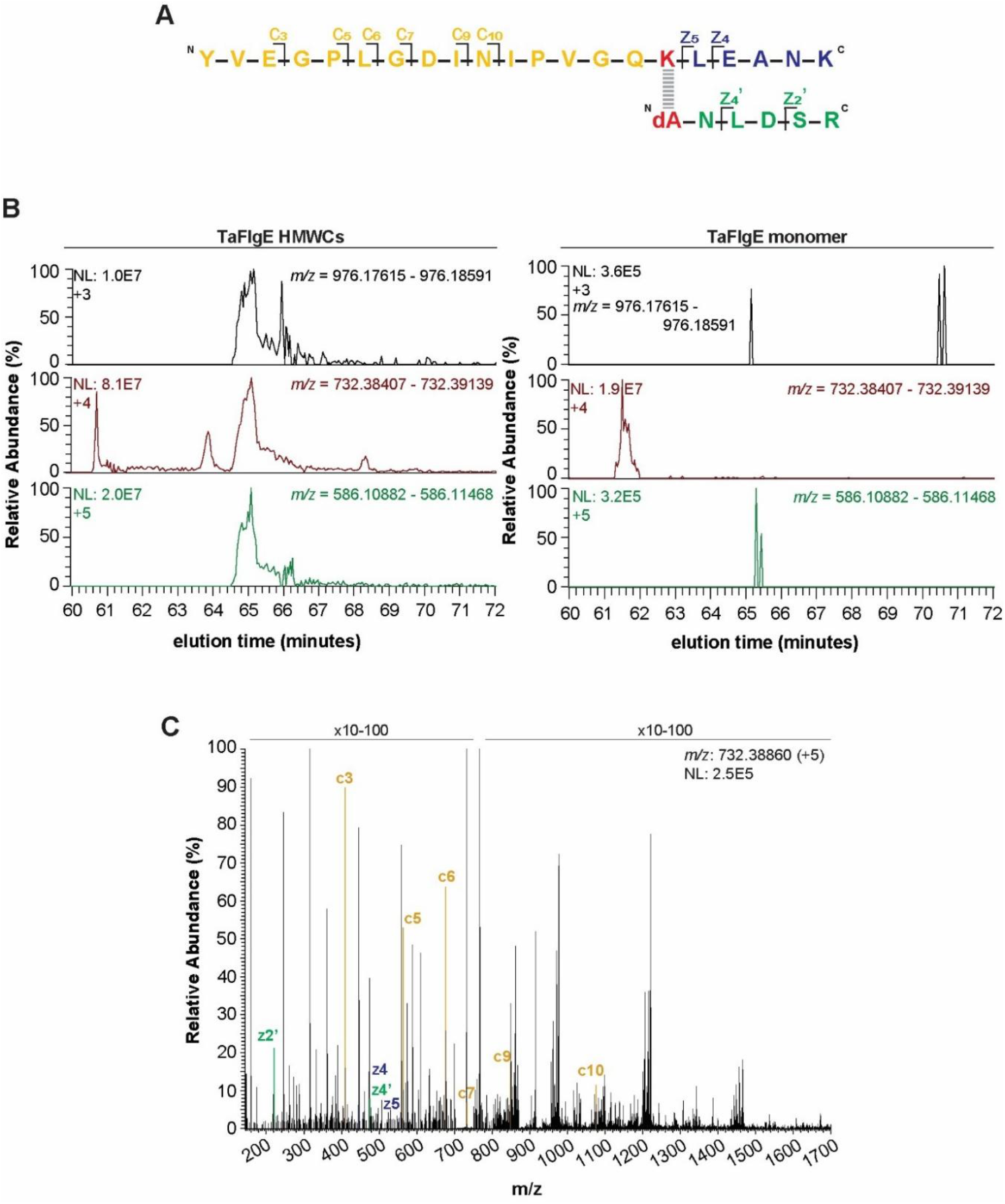
Identification of additional *T. acidaminovorans* FlgE Lal crosslink via LC-MS/MS. A) Primary structure of secondary Ta FlgE Lal crosslinked peptide with c- and z-ions labeled. B) XICs of Lal crosslinked peptide m/z ranges in the [M+3H]^3+^ to [M+5H]^5+^ states from the HMWC (left) and monomer (right) datasets. C) ETD MS/MS fragmentation spectrum of [M+5H]^5+^ parent Lal peptide with c- and z-ions labeled as shown in (A).

**Figure S4:**
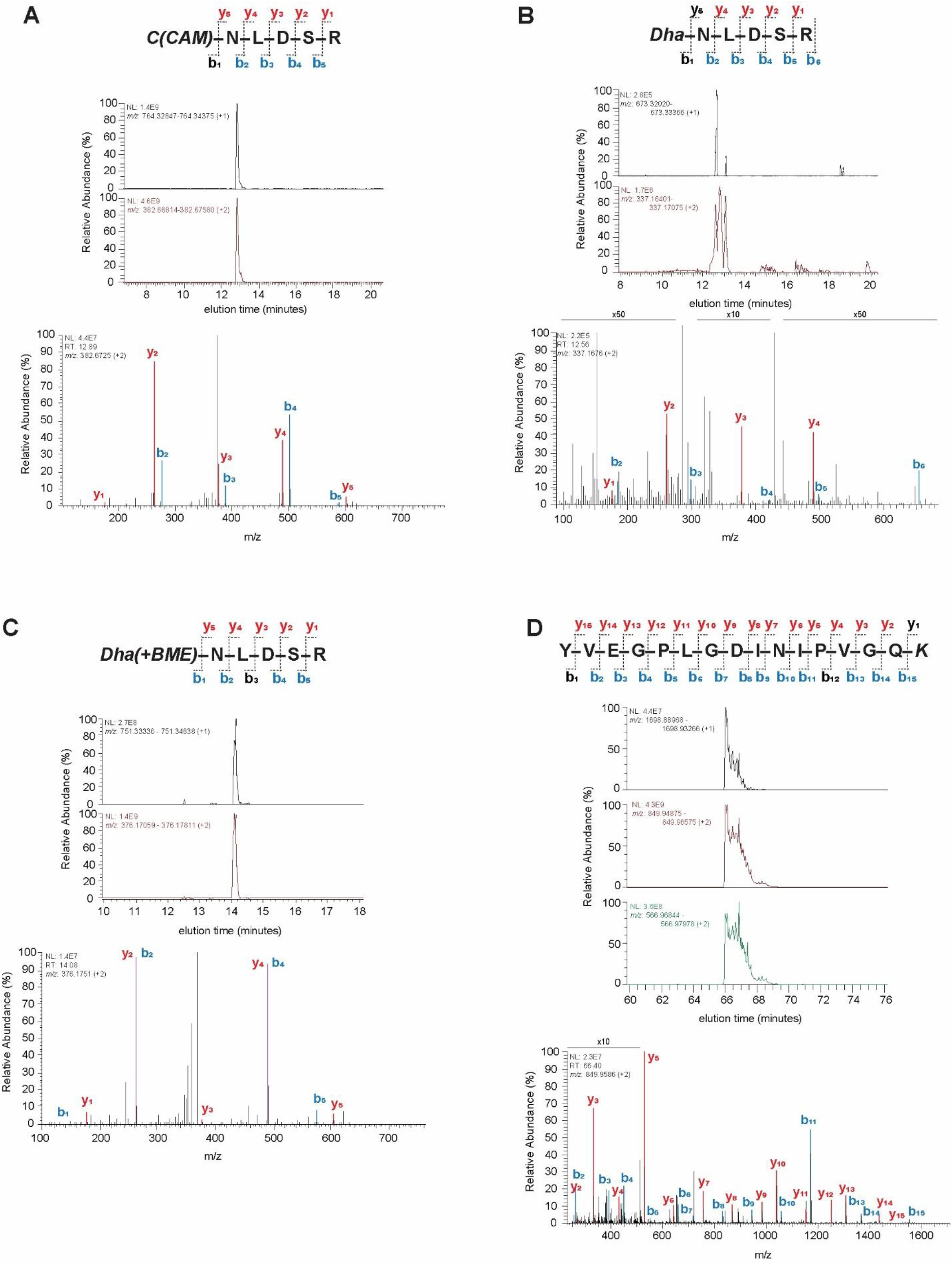
Identification of *T. acidaminovorans* Lal crosslinked parent and intermediate peptides via LC-MS/MS. (top) Primary structure, XICs (middle) and (bottom) CID MS/MS fragmentation spectrum of Lal crosslinking parent peptides: (A) C(CAM)NLDSR, (B) DHA-NLDSR, (C) DHA(BME)-NLDSR and (D) YVEGPLGDINIPVGQK.

**Figure S5:**
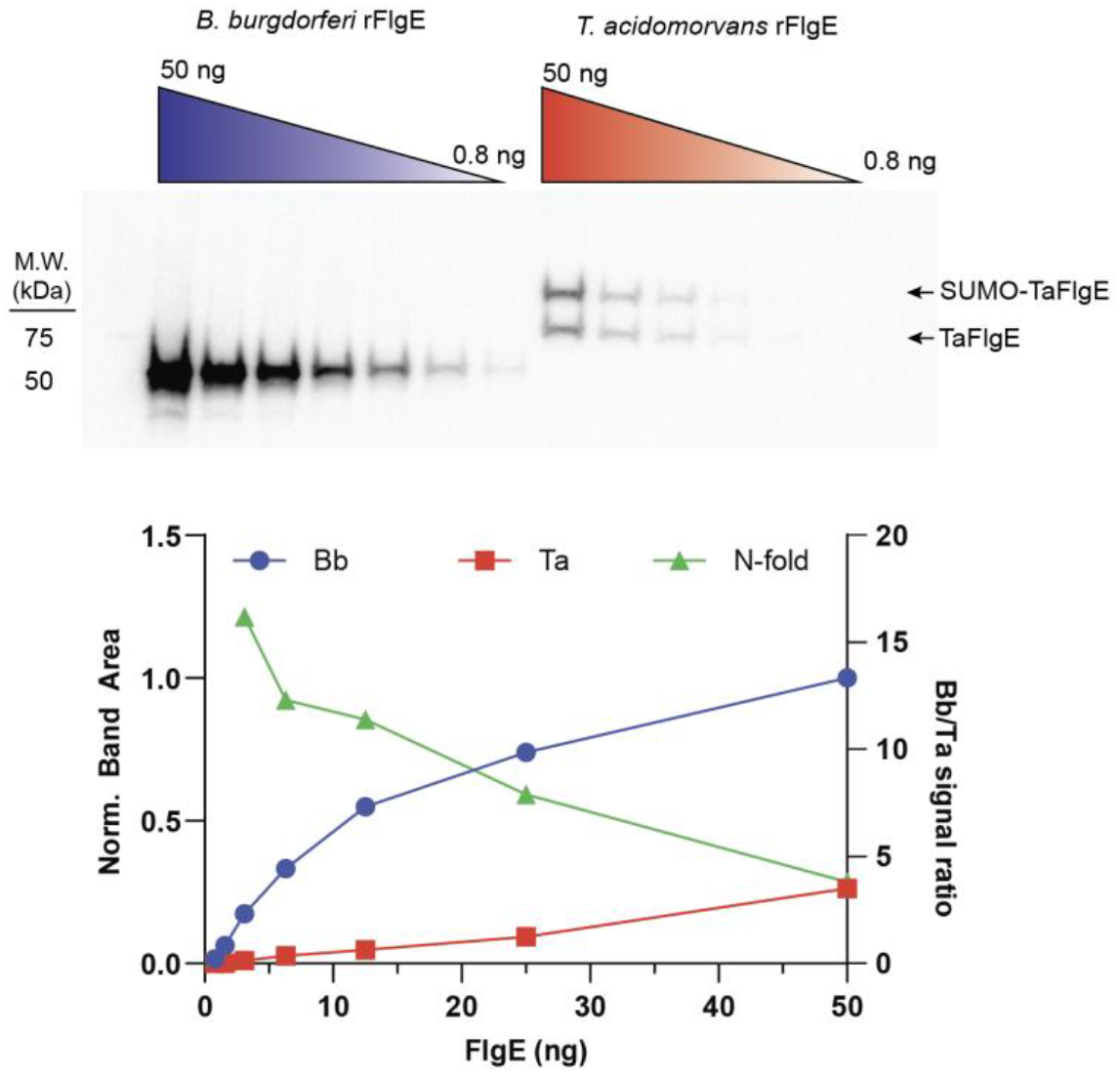
Comparision of anti-FlgE antibody binding to *B. burgdorferi* and *T. acidaminovorans* FlgE. (top) Western blot of decreasing amounts of Bb and Ta FlgE. Notably, our SUMO-TEV-TaFlgE construct failed to be cleaved by TEV fully, so both bands were quantified. (bottom) Quantification of both the Bb (blue, spheres) and Ta (red, squares) FlgE band area normalized to 50 ng Bb FlgE. The N-fold data (green, triangles) was quantified by taking the Bb/Ta FlgE band area ratio.

**Figure S6:**
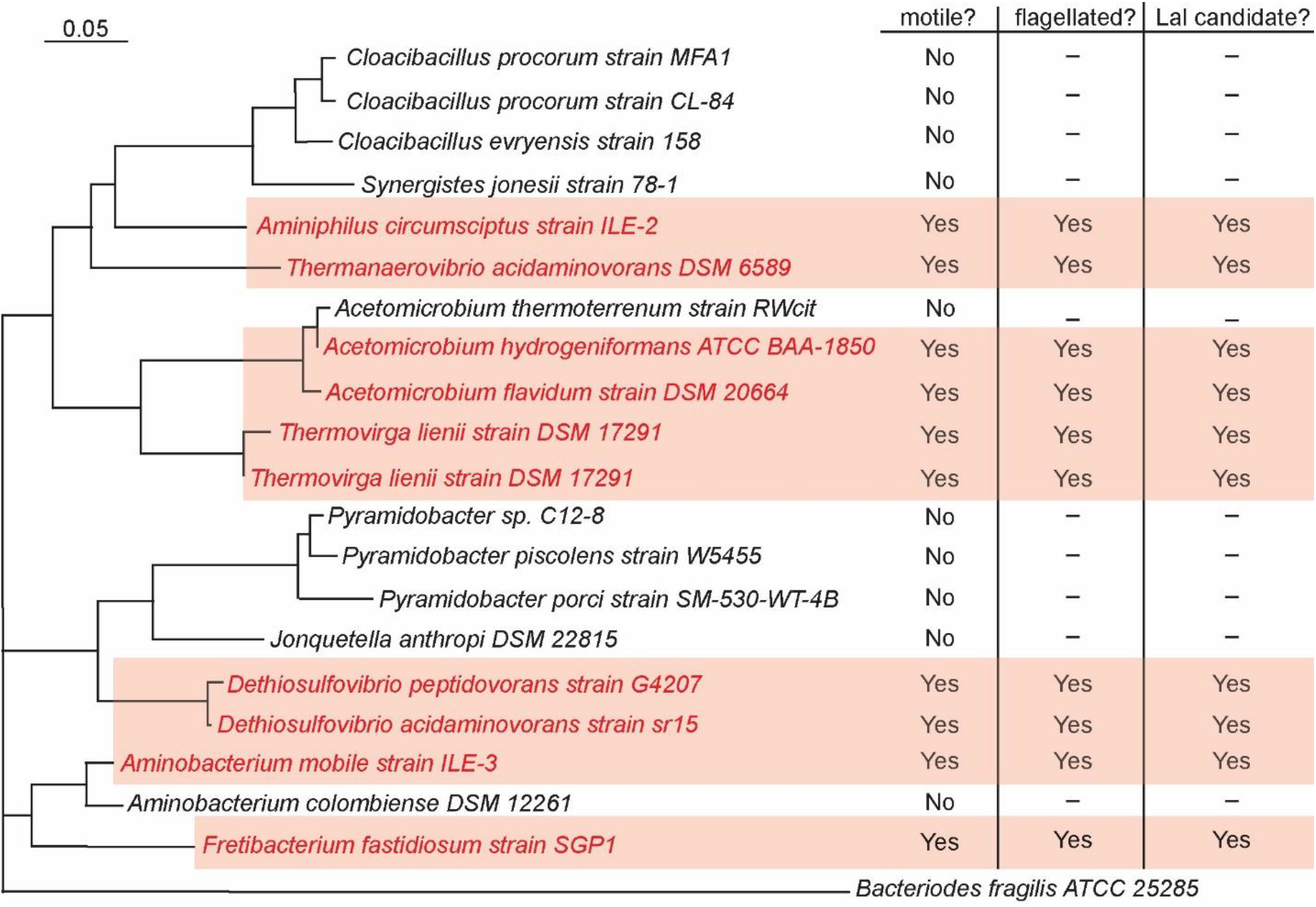
Taxonomic tree of members of the Syngeristota phylum. Taxonomic tree of members of the Syngeristota phylum adapted from McSweeney *et al.,*^50^ generated from near full length 16S rRNA gene sequences. Species that are motile, flagellated and were filtered out as possible Lal crosslinking candidates are shaded in red.

**Figure S7:**
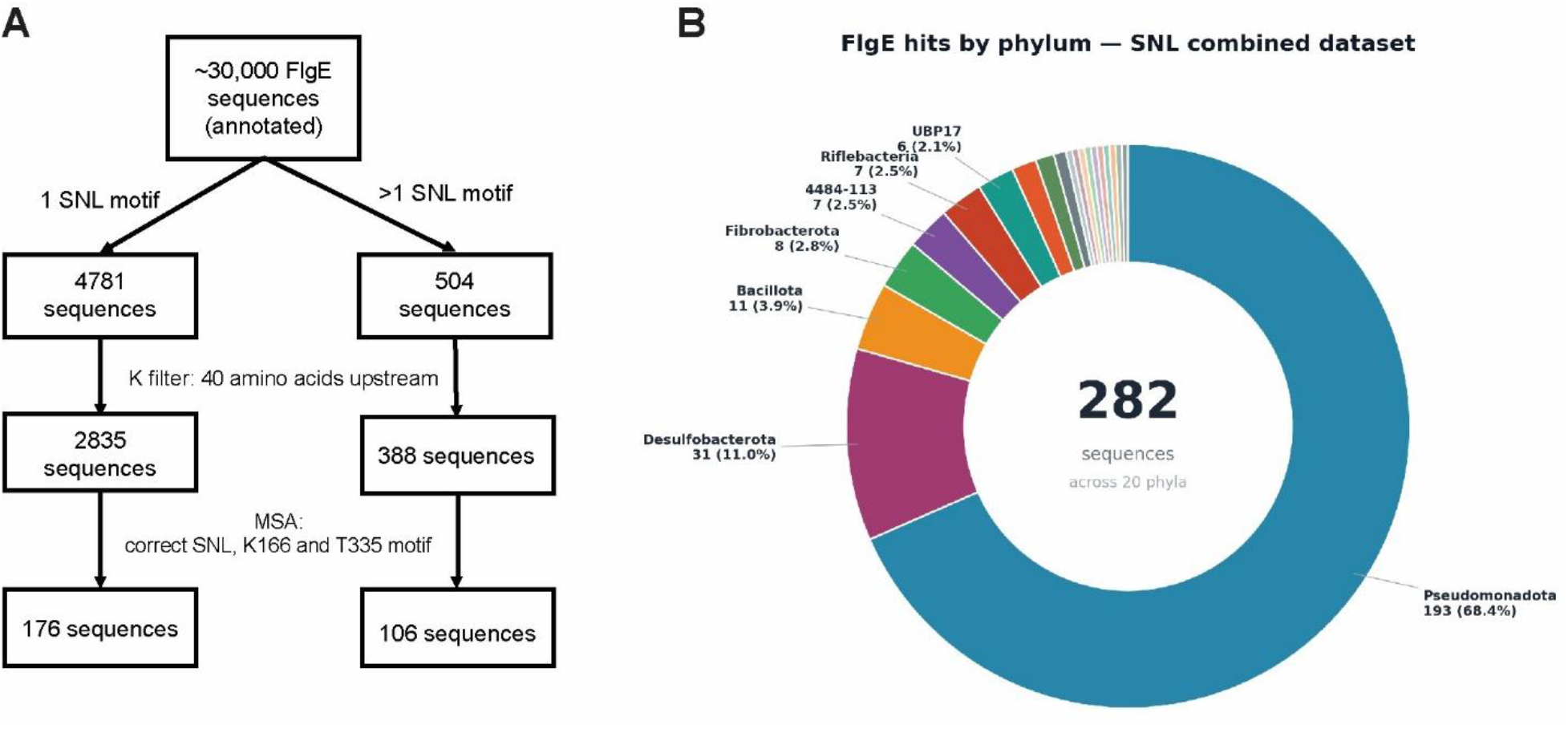
Overview of SNL bioinformatic screen to identify additional Lal candidate species. A) FlgE sequence search strategy using a motif containing K166, SNL, and T335 (Li numbering). Sequences containing one (left) or two (right) SNL motifs were excluded if they lacked a lysine within the 40-amino-acid region upstream of the SNL motif. Candidate sequences were subsequently aligned, and the presence of the catalytic lysine and threonine residues was used to confirm each sequence as a potential Lal candidate. B) Overview of the taxonomic organization of SNL-containing hits identified in the search.

